# Multicellular, fluid flow-inclusive hepatic *in vitro* models using NANOSTACKS^TM^: a human-relevant model for drug response prediction

**DOI:** 10.1101/2024.08.12.607396

**Authors:** Abdullah Talari, Raffaello Sbordoni, Valmira Hoti, Imran I. Patel, Francis L. Martin, Ahtasham Raza, Valon Llabjani

**Affiliations:** REVIVOCELL Limited, Sci-Tech Daresbury, Keckwick Lane, Daresbury, Warrington, WA4 4AD, UK

**Keywords:** NANOSTACKS™, Drug-induced liver injury (DILI), hepatic models, complex *in vitro* models

## Abstract

Drug-induced liver injury (DILI) continues to be the leading cause of drug attrition during clinical trials as well as the number one cause of post-market drug withdrawal due to the limited predictive accuracy of preclinical animal and conventional *in vitro* models. In this study, the NANOSTACKS^TM^ platform was introduced as a novel *in vitro* tool to build *in vivo*-relevant organ models for predicting drug responses. In particular, hepatic models including monocultures of primary human hepatocytes (PHH), tricultures of PHH, human stellate cells (HSC) and human liver endothelial cells (LECs), and tetracultures of PHH, HSC, LECs and human Kupffer cells (KC) were developed under static and fluid flow-inclusive conditions. All hepatic models were characterised by assessing albumin, urea, CYP3A4 and ATP production. In addition, the preclinical DILI screening potential of the fluid flow-inclusive monoculture and triculture models were assessed by testing the hepatotoxicity of Zileuton, Buspirone and Cyclophosphamide. NANOSTACKS™ represents a promising tool for the development of complex *in vitro* models.

## Introduction

Drug-induced liver injury (DILI) represents a relatively rare yet significant source of acute and chronic liver disease (1). It is one of the most common causes of post-market drug withdrawal (2). The detection of hepatotoxicity usually occurs during clinical trials or after a product has been released to the market, leading to elevated hazards for clinical trial participants and imposing huge financial strains on drug development research (3). One of the reasons underlying failures in drug development arises from the limited predictive accuracy of preclinical models (4), which involve animal models and conventional *in vitro* models.

Animal models are used for assessing DILI and evaluating pharmacokinetics despite their inter-species differences with humans with regards to physiology, genetics and drug metabolism (5). A comprehensive large-scale study comparing the effectiveness of animal models in detecting DILI in humans indicated poor predictive performance (6). Among the 150 hepatotoxins studied, both rodent and non-rodent models were only able to detect 50% of the human hepatotoxic events associated with these drugs (6). To address this issue, *in vitro* models replicating aspects of human hepatic biology could be utilised.

Most liver *in vitro* models adopted to screen DILI toxicity are typically based on 2D liver monoculture cell models. Nevertheless, these models are primarily constrained by their lack of crosstalk between different cell types, inconsistent findings with regards to the prediction of hepatotoxicity, and lack of tissue-like organization; such factors are essential for establishing a liver model that accurately mimics physiological conditions (7). Recently, the U.S. Food and Drug Administration (FDA) Modernization Act 2.0 has highlighted the need for alternatives to animal testing and to traditional *in vitro* models, such as 3D *in vitro* models based on the use of organoids and microphysiological systems (8). In the context of hepatotoxicity assessment, alternatives to traditional *in vitro* models include sandwich-cultured hepatic cells, whole organ explants, precision-cut tissue slices, tumour tissue explants, hepatic spheroids, organoids, and liver models developed using microfluidic systems (9–13). In comparison to traditional *in vitro* models, these advanced models can simulate and predict cellular behaviour and therapeutic responses with higher reliability (14–15).

The type of cells included in the *in vitro* model also plays a key role in determining its capability to predict hepatotoxicity. The human liver consists of two main types of cells: hepatocytes and non-parenchymal cells (NPC). NPCs include liver endothelial cells, stellate cells and Kupffer cells (16). Primary human hepatocytes (PHH) are considered the gold standard for studying liver function, liver diseases, drug targets and long-term DILI (17–18).

In the context of preclinical *in vitro* screening models, preserving the metabolic function of PHH over an extended period of cell culture is a significant focus. One approach to enhance the metabolic activity of PHH is to place them in coculture with NPC (16). Hepatocytes cocultured with stellate cells display improved physiological and metabolic functions, and their hepatic function is better maintained (19). For instance, PHH co-cultured with stellate cells exhibit a more stable liver phenotype compared to monocultures (20–22). Similarly, the coculture of hepatocytes with liver endothelial cells in a microfabricated perfusion reactor resulted in the formation of endothelial network structures and high retention of hepatocellular function compared to monocultures (23). Various other cellular combinations involving PHH have been investigated, including the use of NIH/3T3 (24, 25), endothelial cells (26, 27), and Kupffer cells (28, 29).

In order to setup a coculture model, a common approach is based on mixing different cell types in a single well (30, 31). The mixture approach offers the advantage of facilitating direct cell-cell contact, allowing the evaluation of interactions mediated by cell adhesion. However, variability in growth rates can result in randomly distributed cell populations, which can significantly differ between each replicate, thus decreasing the reproducibility of the model. Furthermore, this method does not replicate the layered microarchitecture of hepatic lobules. Another approach is constituted by the sandwich culture, based on the culture of hepatic cells in different layers (32, 33). This method models the natural layering of liver cells whilst maintaining constant the ratios between the cell number of different cell types included in the model. However, the sandwich method is time-consuming, challenging to reproduce, and labour-intensive (34). Human liver microphysiological systems (MPS) have the potential to address the limitations of current coculture models by utilizing engineering and design principles that more accurately replicate human liver physiology in miniatured systems. These advanced models may incorporate various sophisticated features such as a multicellular environment, 3D architecture and exposure to fluid flow. However, MPS-based models can be complicated to develop and use, therefore reducing their applicability in the context of DILI screening.

To address issues associated with currently used models, in this this work NANOSTACKS^TM^ (NS), a novel user-friendly, imaging- and fluid flow-compatible platform, was used for the assembly of complex hepatic cocultures in a 24-well plate format. In particular, hepatic models based on monocultures of PHH, tricultures of PHH, human stellate cells (HSC) and human liver endothelial cells (LECs), and tetracultures of PHH, HSC, LECs, and human Kupffer cells (KC) on NS were developed. The models were characterised with regards to parameters associated with hepatic function, in absence or presence of fluid flow. Finally, three different compounds (Zileuton, Buspirone and Cyclophosphamide) were tested on the fluid flow-inclusive triculture and monoculture hepatic models to assess their reliability for preclinical DILI screening.

## Methods

### Cell culture

Cryopreserved primary human hepatocytes (PHH) (Lot 2211419-01), primary human liver endothelial cells (LECs) (Lot 2211419p0), primary human stellate cells (HSC) (Lot 2216631p0) and primary human Kupffer cells (KC) (Lot 2211419) were purchased from LifeNet Health LifeSciences. All primary cells were cultured in accordance with the protocols specified by the vendor. Before seeding on NS, both LECs and HSC were expanded on rat collagen type I (Gibco^TM^)-coated T-25 flasks (Fisher Scientific). LECs were expanded using LECs complete medium, composed of Lonza EBM-2 and Lonza EGM-2, whilst HSC were expanded using HSC complete medium, composed of DMEM (Gibco^TM^), 10% Fetal Bovine Serum (FBS) (Sigma) and 1% Pen/Strep (Sigma). PHH and KC were thawed according to the vendor’s protocols on the day of seeding on NS. PHH thawing medium, plating medium and maintenance medium were provided by LifeNet Health LifeSciences. KC complete medium, which was used to culture KC on NS before initiating the tetraculture, was composed of RPMI 1640 (Gibco^TM^), 10% Fetal Bovine Serum (FBS) (Sigma) and 1% Pen/Strep (Sigma).

### NANOSTACKS^TM^ (NS) design

NS enable 3D cell culture through their stackable design within a standard SBS 24-well plate format (Figure 1). In particular, up to four NS can be stacked in a single well, and therefore up to four different cell types can be cocultured within the same *in vitro* model. Cell culture medium can diffuse across the gaps between each NS. Additionally, each NS includes a porous membrane, with a pore size of 0.4 µm, increasing nutrients diffusion towards cells. Additionally, the porous membrane of NS is transparent, thus allowing live imaging without disrupting the multilayered structure. To avoid material absorption of pharmaceutical compounds, the body of NS are composed of polycarbonate whilst the membrane is composed of polyester. In order to induce fluid flow on the NS, the 24-well plates including the devices can be placed on an orbital shaker.

**Figure 1.**
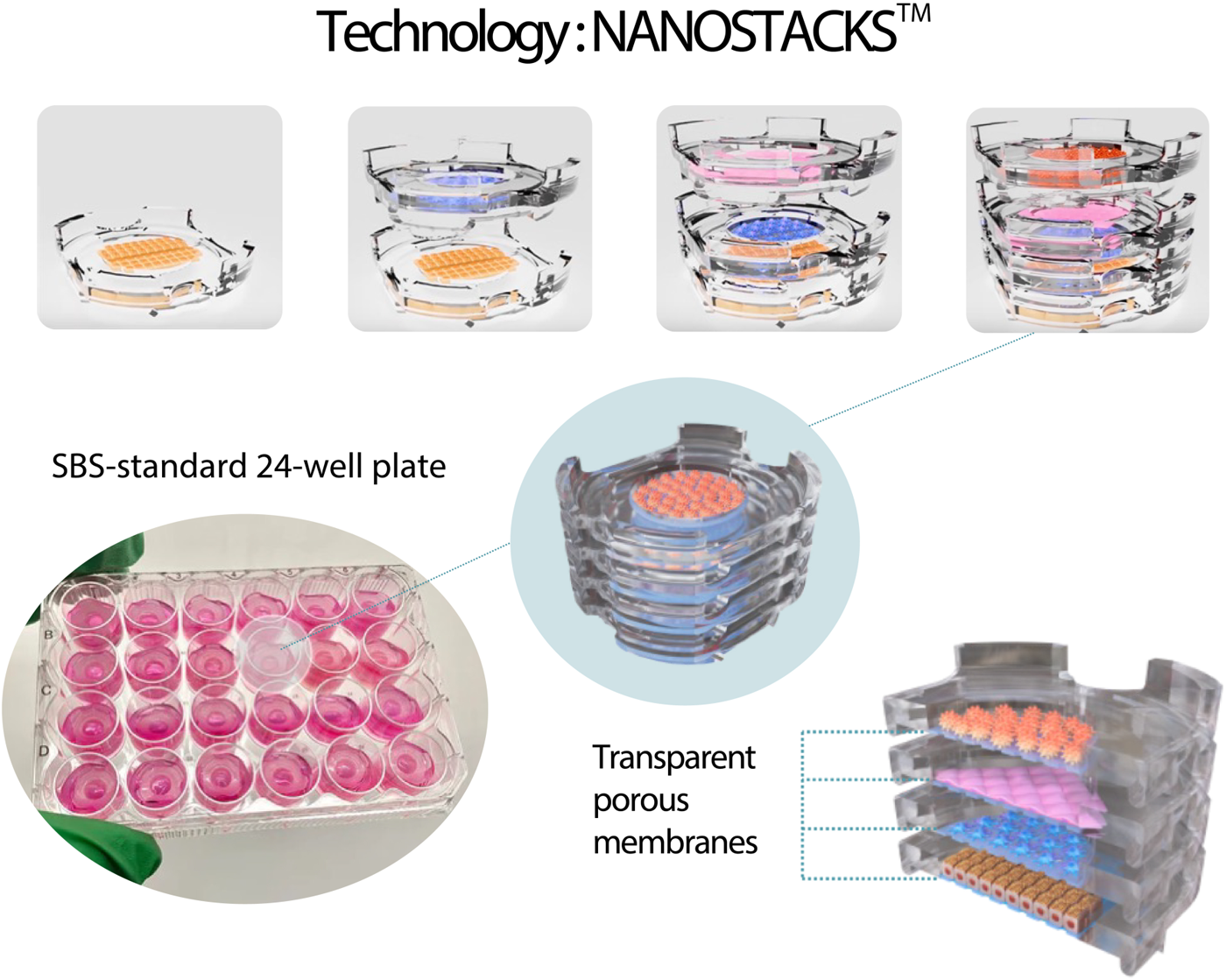
Graphic representation of the NS platform. Up to four cell-seeded NS can be stacked on top of each other (top), forming a complex coculture (centre) that can be housed into wells of a SBS-standard 24-well plate (bottom-left). Each NS is made of a polycarbonate body and a transparent porous PET membrane with a pore size of 0.4 μm (bottom right). Fluid flow can be induced by placing the 24-well plate including the NS on an orbital shaker.

### Human liver modelling on the NANOSTACKS™ (NS) platform

Three types of human liver models were developed using NS. In particular, the models were a monoculture model (PHH), a triculture model (PHH + LECs + HSC) and a tetraculture model (PHH + LECs + HSC + KC). All the models were developed with and without the inclusion of fluid flow. Throughout the experiment, NS were kept in wells of 24-well plates. All NS were coated with 10 μg/cm^2^ rat collagen type I at RT and then washed thrice using phosphate buffered saline (PBS). PHH, LECs, HSC and KC were seeded at a 3:1:1:1 proportion respectively to the seeding density of the individual cell type. In particular, a cell seeding suspension volume of 70 μL was decanted on the top surface of the cell culture-treated membrane on each NS, and cells were incubated at 37°C and at 5% CO_2_ in a humidified incubator for 2 h to allow cell attachment. PBS was added to empty wells of the well-plate to prevent evaporation of the cell suspension droplets. Then, 1430 μL of medium was added to the seeded NS to reach the working volume of 1.5 mL and subsequently the plates were left in a cell culture incubator for 24 h. Each cell type was seeded using its respective complete medium. On Day -1, each NPC (LECs, HSC, and KC) was seeded at a seeding density of 16.6 x 10^3^ cells/mL whereas on Day 0 PHH were seeded at a seeding density of 50 x 10^3^ cells/mL. On Day 1, the *in vitro* models were assembled by stacking the cell-seeded NS in order to combine the different cell types into monoculture, triculture and tetraculture models (Fig. 2A). In particular, in the monoculture model one PHH-seeded NS was placed in each well. In the triculture model, one PHH-seeded NS was placed in the bottom of each well, and one HSC-seeded, one LECs-seeded were placed on top. In the tetraculture model, one PHH-seeded NS was placed in the bottom of each well, and one HSC-seeded, one LECs-seeded, and one KC-seeded NS were placed on top of the PHH-seeded NS. Additionally, for monoculture models, three membrane-free, cell-free NS were added on top of the cell-seeded NS, whilst for triculture models, one membrane-free, cell-free NS was added on top of the three cell-seeded NS, in order to maintain the same height of cell culture medium in the wells of all hepatic models. Once the models were assembled, PHH culture medium was used to maintain the cultures, with medium changes occurring every 2 days. On day 1, each model was assigned either to a static condition or to a fluid flow condition, the latter entailing the inclusion of fluid flow by placing the 24-well plate including the NS on an orbital shaker (TOS-3530CO2; Munro Scientific) set at 90 RPM. Monocultures and tricultures were maintained in culture for 31 days, whilst tetracultures were maintained for 26 days. All models underwent characterization based on cell viability, albumin, urea, and CYP3A4 metabolic activity measurements. DILI toxicity screening was conducted exclusively on the monoculture and triculture models. Each assay was performed on n = 3 wells per timepoint in both static and fluid flow conditions.

**Figure 2.**
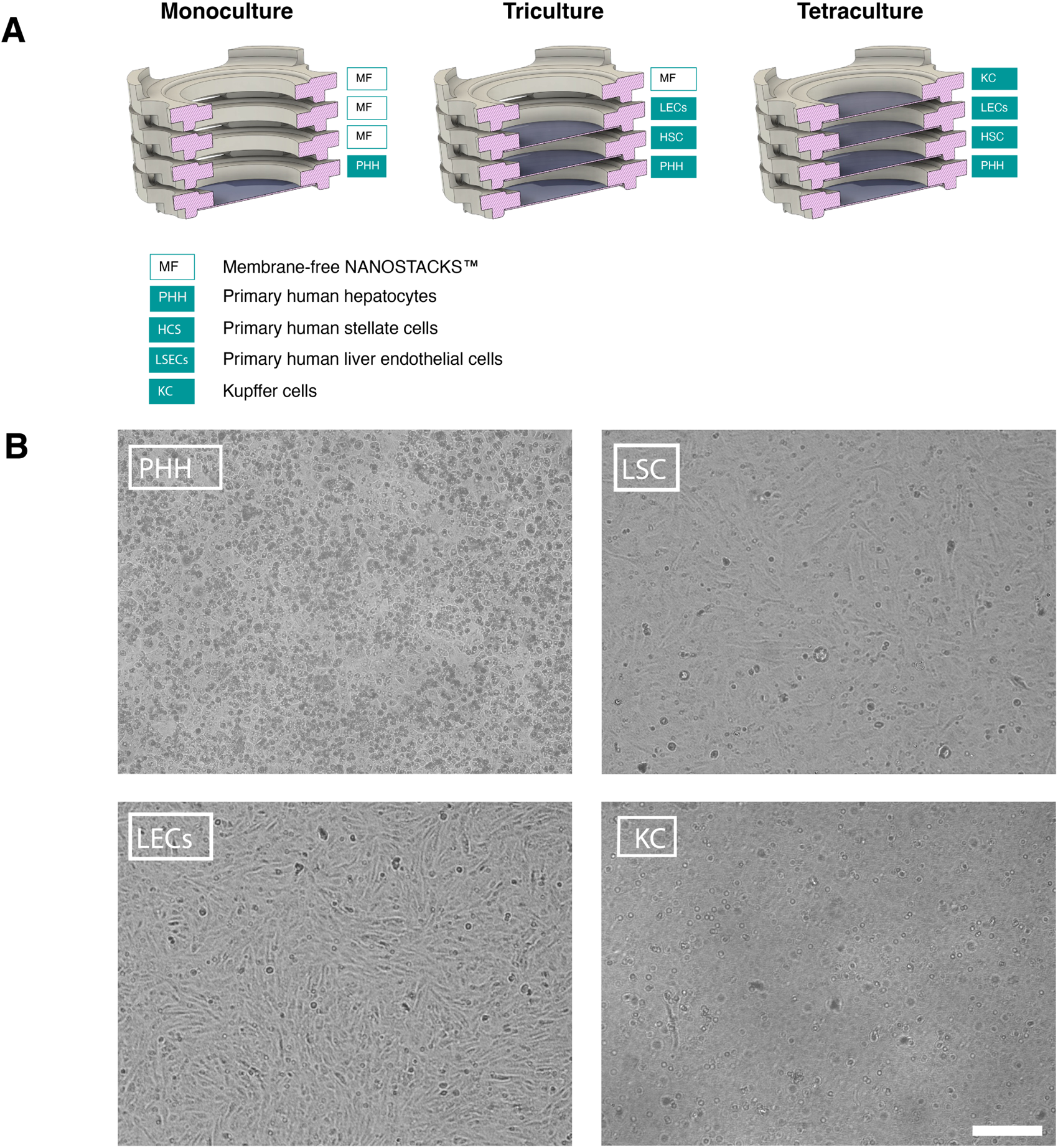
**A:** Schematic representation of the experimental setup. **B:** Representative widefield image of primary human hepatocytes (PHH), primary human stellate cells (HSC), primary human liver endothelial cells (LECs), and primary human Kupffer cells (KC) on NS, acquired on day 2. Magnification: 10X. Scale bar: 250 μm

### Cell viability

The CellTiter-GLO assay (CellTiter-Glo® Luminescent Cell Viability Assay G7571, Promega), which measures intracellular ATP content as a biomarker of cell viability, was performed according to the protocol provided by the vendor, with the following modifications: cell-seeded NS were moved to wells of a 24-well plate including 350 μL of medium, and then 350 μL CellTiter-GLO reagent was added onto each well to obtain a 1 : 1 dilution. The luminescence of each well was read by a Synergy H1 Microplate reader (Fisher Scientific) and analysed with the GEN5 software (BioTek; version 2.05). The CellTiter-GLO assay was performed on n = 3 wells per timepoint on days 2, 4, 7, 11, 14, 20, 26 and 31.

### Cytochrome P450 assay

CYP3A4 expression in PHH was measured with a P450-GLO assay (V9001 Luciferin-IPA, Promega). The P450-Glo™ assay technology provides a rapid, high-throughput method for assessing cytochrome P450 (CYP) activity by measuring the conversion of inactive D-luciferin derivatives to an active form. The emitted light intensity is directly proportional to the CYP enzyme activity. The assay was performed on day 2, 4, 7, 11, 14, 20, 26 and 31 according to the protocol provided by the vendor, on the same NS used for viability analysis. The assay was performed on n = 3 wells per timepoint.

### Albumin Assay

The albumin produced by cells within the liver models was quantified on days 2, 4, 7, 11, 14, and 26 using sandwich ELISA kits (Albumin ab179887, Abcam). In particular, the supernatant obtained from each well was preserved at −80 °C, then thawed overnight at 4 °C. Subsequently, the supernatant samples were diluted 50-fold with the buffer provided by the vendor and the assay was performed following vendor’s instructions. In the last step of the assay, absorbance at 450 nm was measured using the Synergy H1 Microplate reader and analysed with the GEN5 software. The assay was performed on n = 3 wells per timepoint.

### Urea Assay

Urea produced by PHH in the liver models was quantified using a urea assay kit (MAK006, Sigma-Aldrich) on days 2, 4, 7, 11, 14, and 26. The supernatant obtained from each well was preserved at −80 °C, and thawed overnight at 4 °C. Subsequently, the supernatant samples were diluted 50-fold with the buffer provided by the vendor. The assay was then performed according to the instructions of the vendor. Finally, absorbance at 570 nm was measured using the Synergy H1 Microplate reader and analysed with the GEN5 software. The assay was performed on n = 3 wells per timepoint.

### Toxicity screening

The dose-response effects of Zileuton, Buspirone hydrochloride and Cyclophosphamide were assessed at concentrations ranging from 1 to 600 times the human C_max_ on monoculture and triculture models. The experiments were conducted on day 4 on monocultures and on day 7 on tricultures models. The compounds were dissolved in DMSO, which was used at concentration of 0.1 % V/V in cell culture medium and was also included as vehicle control. The models were treated with the compounds every day for 7 days. Total cytotoxicity of the compounds was then measured using the CellTiter-Glo® Luminescent Cell Viability Assay (G7571, Promega), as previously described. Drug treatments were performed in n = 3 wells per compound and concentration.

### Fluid dynamics modelling on NANOSTACKS^TM^

A computational fluid dynamics (CFD) model was developed using the software ANSYS-CFX (ANSYS Inc.) to model the shear stress exerted on NS placed into wells of a 24-well plate in an orbital shaker set at 90 RPM. In particular, CFD modelling was performed on a well including one NS inclusive of membrane (bottom of the well) and three NS without membranes, on three NS inclusive of membranes (bottom of the well) and one NS without membrane, on four NS including membranes, and on an NS-free well. The orbital diameter of the orbital shaker was 19 mm and the volume of cell culture medium was 1.5 mL in all configurations. The medium was modelled as an incompressible and Newtonian fluid with a dynamic viscosity (µ) of 0.7 mPa·s at 37 °C and medium density ρ = 1000 kg/m^3^ as described by Driessen et al. (35).

### Statistical analysis

For each timepoint associated with ATP, CYP3A4, albumin and urea production, unpaired Student T-tests were performed using the software Prism (GraphPad; version 10.2.3) to analyse the differences between the static and fluid flow conditions.

A p-value < 0.05 was assumed to indicate a statistically significant difference, indicated by asterisks in the graphs (*: p < 0.5; **: p < 0.01; ***: p < 0.001; ****: p < 0.0001). Values are reported in the graphs as mean ± SEM.

## Results

In this work, NS-based monocultures (PHH), tricultures (PHH-HSC-LECs) and tetracultures (PHH-HSC-LECs-KC) liver models were developed under static and fluid flow conditions, the latter induced by placing the 24-well plate including the NS onto a orbital shaker. To quantify the shear stress exerted on the cell culture surface of the inserts in monoculture and coculture configurations, a computational fluid dynamics (CFD) analysis was performed (Fig. 3). The shear stress was highest on the membrane included in NS occupying the position closest to the air-liquid interface in cocultures (Fig. 3.C-D), comparatively to other NS membranes positioned closer to the bottom of the well (Fig. 3B-D). Additionally, the shear stress profile of the NS membrane associated with the monoculture condition (Fig. 3B) was found to be more spatially uniform than the shear stress profile related to the bottom surface of an empty well (Fig. 3A).

**Figure 3.**
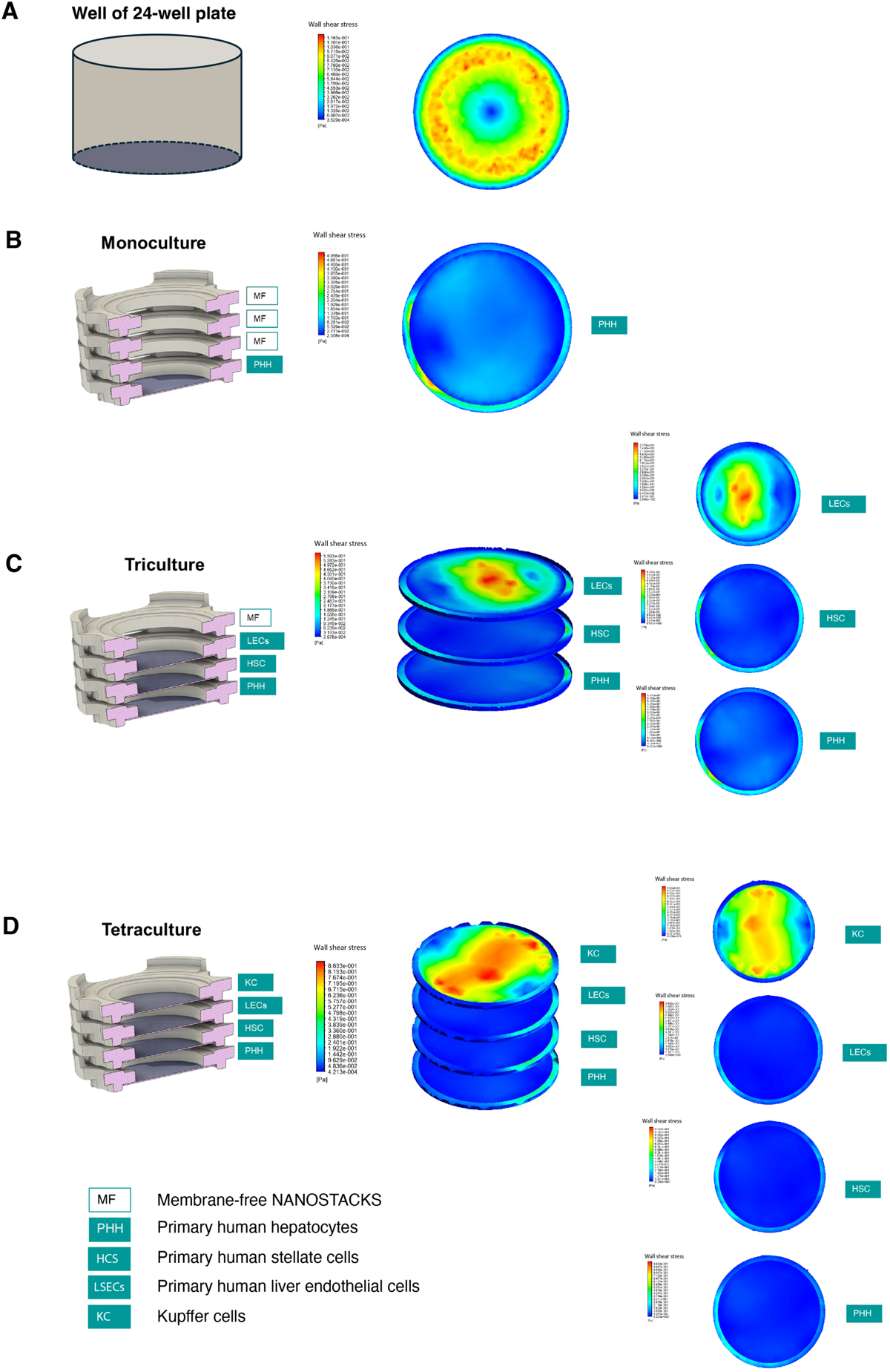
CFD modelling of shear stress (Pa) induced by an orbital shaker set at 90 rpm, acting on an empty well of a 24-well plate and on NS membranes included in the well. **A:** Bottom surface of NS-free well. **B:** Membrane of NS placed on bottom of the well, under three membrane-free NS. **C:** Membranes of three NS placed on the bottom of the well, under one membrane-free NS. **D:** Membranes of four NS placed inside the well.

The NS-based liver models were also characterised in both static and fluid flow conditions by quantifying ATP and CYP3A4 production by PHH, in addition to albumin and urea synthesis. With regards to the monoculture models, the characterisation data is summarised in Figure 4. In particular, ATP production was maintained for 31 days (Fig. 4A), indicating cell viability throughout the entire experiment, with no statistically significant difference between the static and fluid flow conditions. CYP3A4 production was also maintained throughout 31 days (Fig. 4B), peaking on day 7 in all conditions, when the CYP3A4 production was higher in the fluid flow condition relatively to the static condition. Albumin production (Fig.4C) was maintained for 26 days in all conditions, peaking on day 7, reaching 40.1 µg/day/10^6^ PHH in the flow condition. With regards to urea production (Fig. 4D), human-relevant levels of urea (>56 µg/million/PHH/day) (39) were maintained for 7 days in the static condition, decreasing to 24.25 µg by day 14. Conversely, when fluid flow was introduced into the model, urea production was above the human level threshold until day 14.

**Figure 4.**
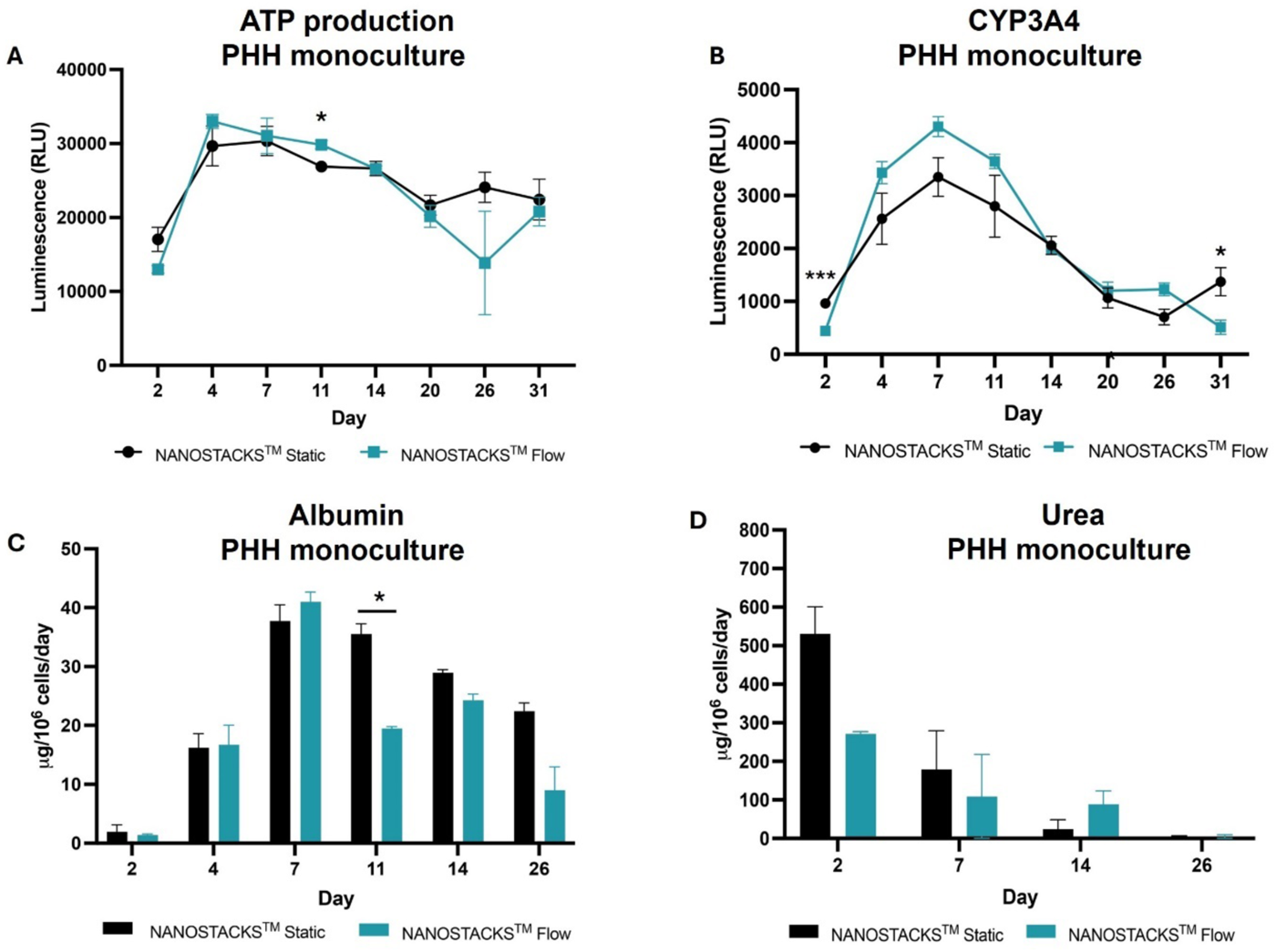
ATP, CYP3A4, albumin and urea production from PHH in monocultures in static (black) and fluid flow (blue) conditions. **A:** ATP synthesis, expressed in relative light units (RLU; y axis) on day 2, 4, 7, 11, 14, 20, 26, 31 (x axis). **B:** CYP3A4 production, expressed in RLU (y axis), on day 2, 4, 7, 11, 14, 20, 26, 31 (x axis). **C:** Albumin production, expressed in µg/10^6^ cells/day (y axis), on day 2, 4, 7, 11, 14, 26 (x axis). **D:** Urea production, expressed in µg/10^6^ cells/day (y axis) on day 2, 7, 14, 26 (x axis). Each datapoint was obtained from n = 3 wells. Data are reported as mean ± SEM.

**Figure 5.**
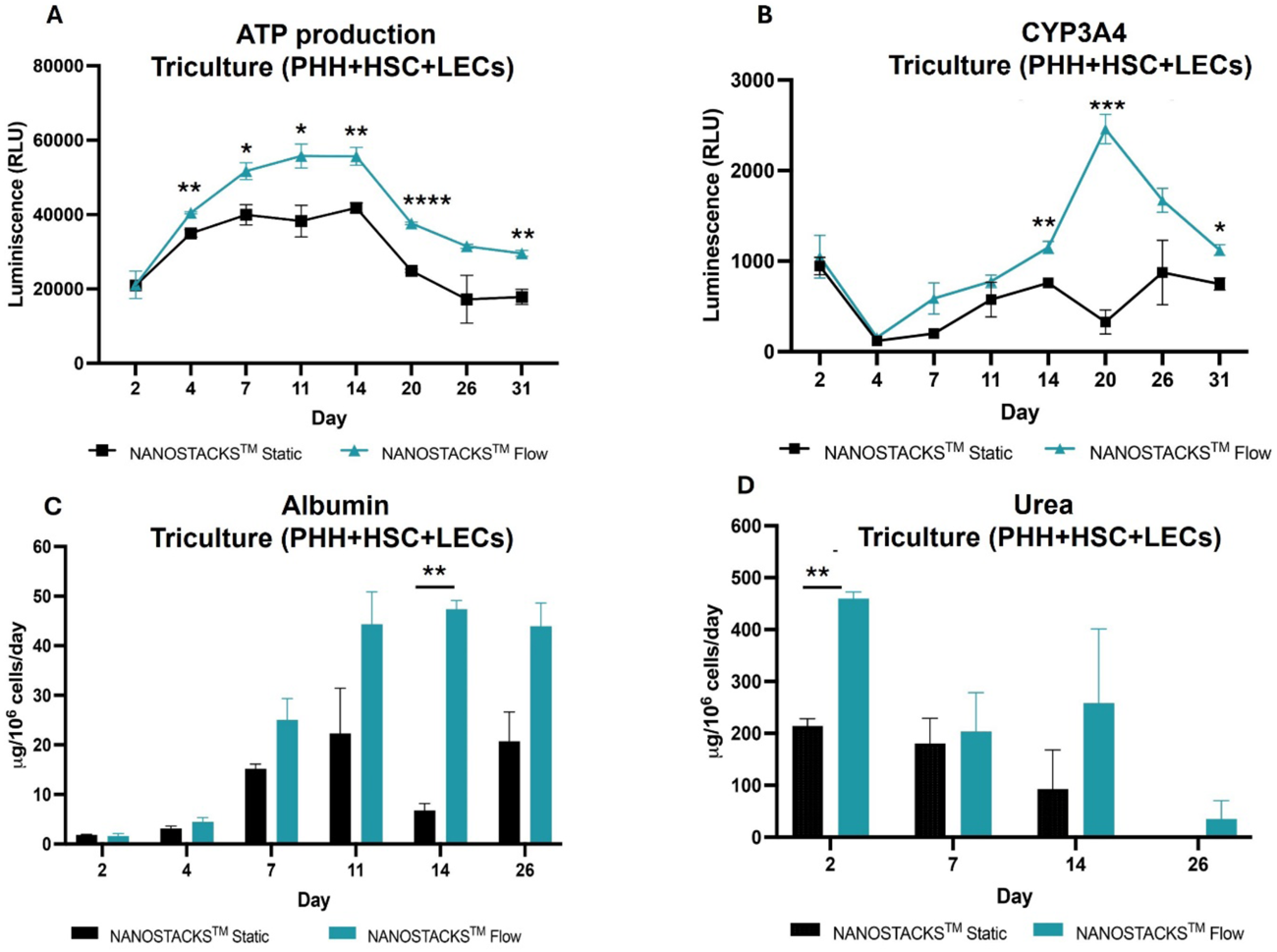
ATP, CYP3A4, albumin and urea production from PHH in tricultures in static (black) and fluid flow (blue) conditions. **A:** ATP synthesis, expressed in RLU (y axis) on day 2, 4, 7, 11, 14, 20, 26, 31 (x axis). **B:** CYP3A4 production, expressed in RLU (y axis), on day 2, 4, 7, 11, 14, 20, 26, 31 (x axis). **C:** Albumin production, expressed in µg/10^6^ cells/day (y axis), on day 2, 4, 7, 11, 14, 26 (x axis). **D:** Urea production, expressed in µg/10^6^ cells/day (y axis) on day 2, 7, 14, 26 (x axis). Each datapoint was obtained from n = 3 wells. Data are reported as mean ± SEM.

PHH in triculture with liver endothelial cells (LECs) and stellate cells grown on NS were viable throughout the entire experiment in both static and fluid flow conditions, as evident from the ATP production sustained for 31 days (Fig. 5A). However, in the fluid flow condition, ATP production was elevated compared to the static condition from day 4. Similarly, CYP3A4 production (Fig. 5B) was also sustained for 31 days. In particular, on day 20 the CYP3A4 production associated with the fluid flow condition was more than seven times higher than the levels reached in the static condition. Albumin production (Fig. 5C) was maintained throughout the entire experiment, and between day 7 and day 26, albumin levels in the fluid flow condition were markedly higher relatively to the static condition. In particular, between day 11 and day 26, albumin production in the fluid flow condition exceeded the human liver *in vivo* output threshold of 43 µg/day/10^6^ PHH (39). With regards to urea production (Fig. 5D), values obtained from both conditions were superior to human threshold levels (>56 µg/million/PHH/day) from day 2 to 14. Additionally, on day 2 in the flow condition, urea production was markedly higher than the production associated with the static condition.

**Figure 6.**
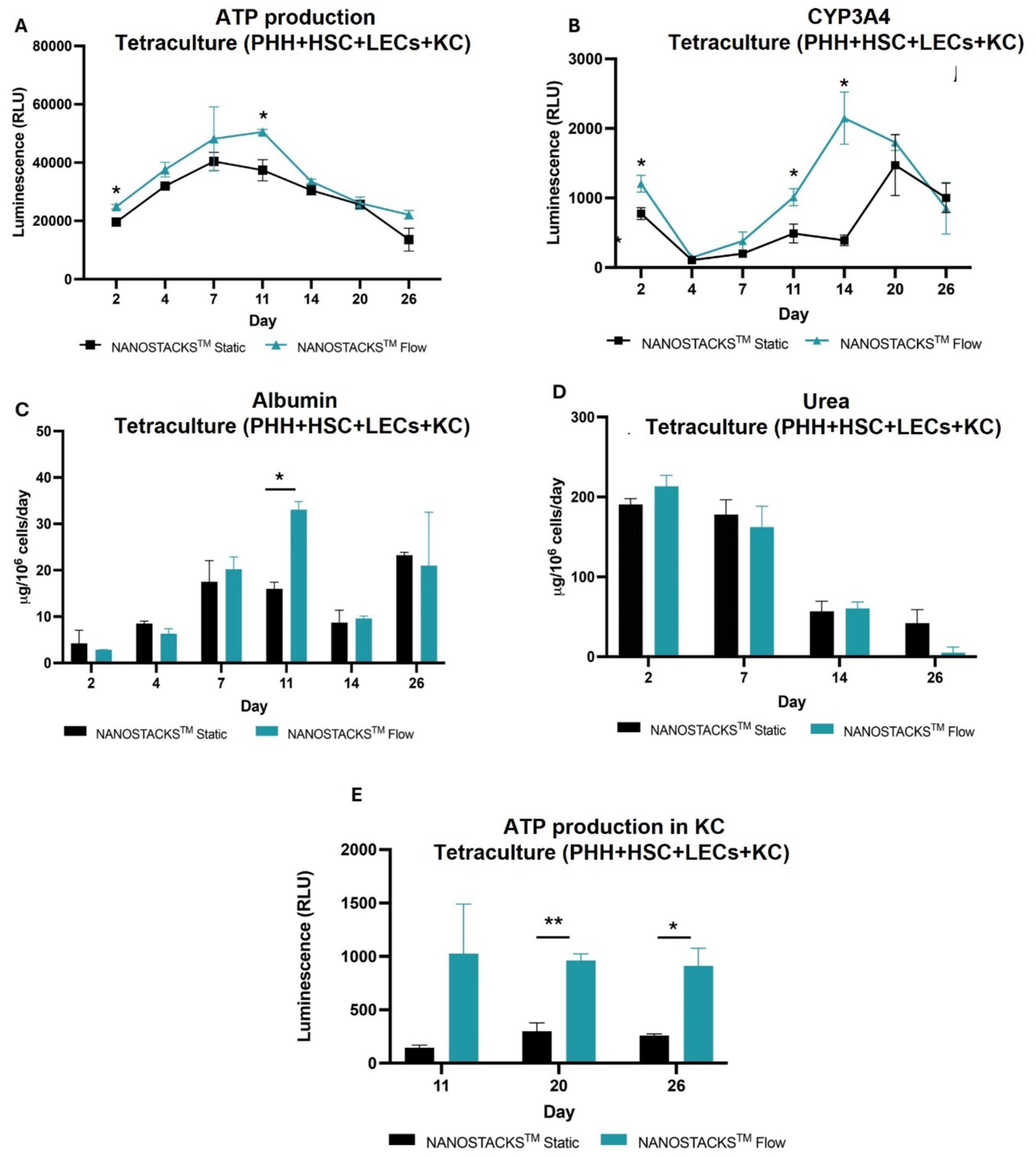
ATP, CYP3A4, albumin and urea production from PHH, and ATP synthesis by Kuppfer cells in tetracultures, in static (black) and fluid flow (blue) conditions. **A:** ATP synthesis by PHH, expressed in RLU (y axis) on day 2, 4, 7, 11, 14, 20, 26 (x axis). **B:** CYP3A4 production, expressed in RLU (y axis), on day 2, 4, 7, 11, 14, 20, 26 (x axis). **C:** Albumin production, expressed in µg/10^6^ cells/day (y axis), on day 2, 4, 7, 11, 14, 26 (x axis). **D:** Urea production, expressed in µg/10^6^ cells/day (y axis) on day 2, 7, 14, 26 (x axis). **E:** ATP synthesis by Kuppfer cells, expressed in RLU (y axis) on day 11, 20 and 26 (x axis). Each datapoint was obtained from n = 3 wells. Data are reported as mean ± SEM.

PHH in tetracultures maintained viability for the entire experiment in both fluid flow and static conditions, as evident from their ATP production (Fig. 6A), which was sustained for 26 days. On day 11, the ATP production associated with the fluid flow condition was markedly higher than the value associated with the static condition. PHH in tetracultures also maintained CYP3A4 production at all timepoints (Fig. 6B). In particular, on day 11 and day 14, CYP3A4 levels associated with the fluid flow condition were higher relatively to the static condition. Additionally, on day 11 albumin production was higher in the fluid flow condition relatively to the static condition (Fig. 6C). Albumin production was maintained in both conditions throughout the entire experiment. Urea production in tetraculture displayed a descending trend in both conditions (Fig. 6D) and was consistently above the human level threshold (>56 µg/million/hepatocytes) for 14 days in both conditions (39). With regards to Kuppfer cells, ATP production was markedly increased in the fluid flow condition relatively to the static condition (Fig. 6E).

**Figure 7.**
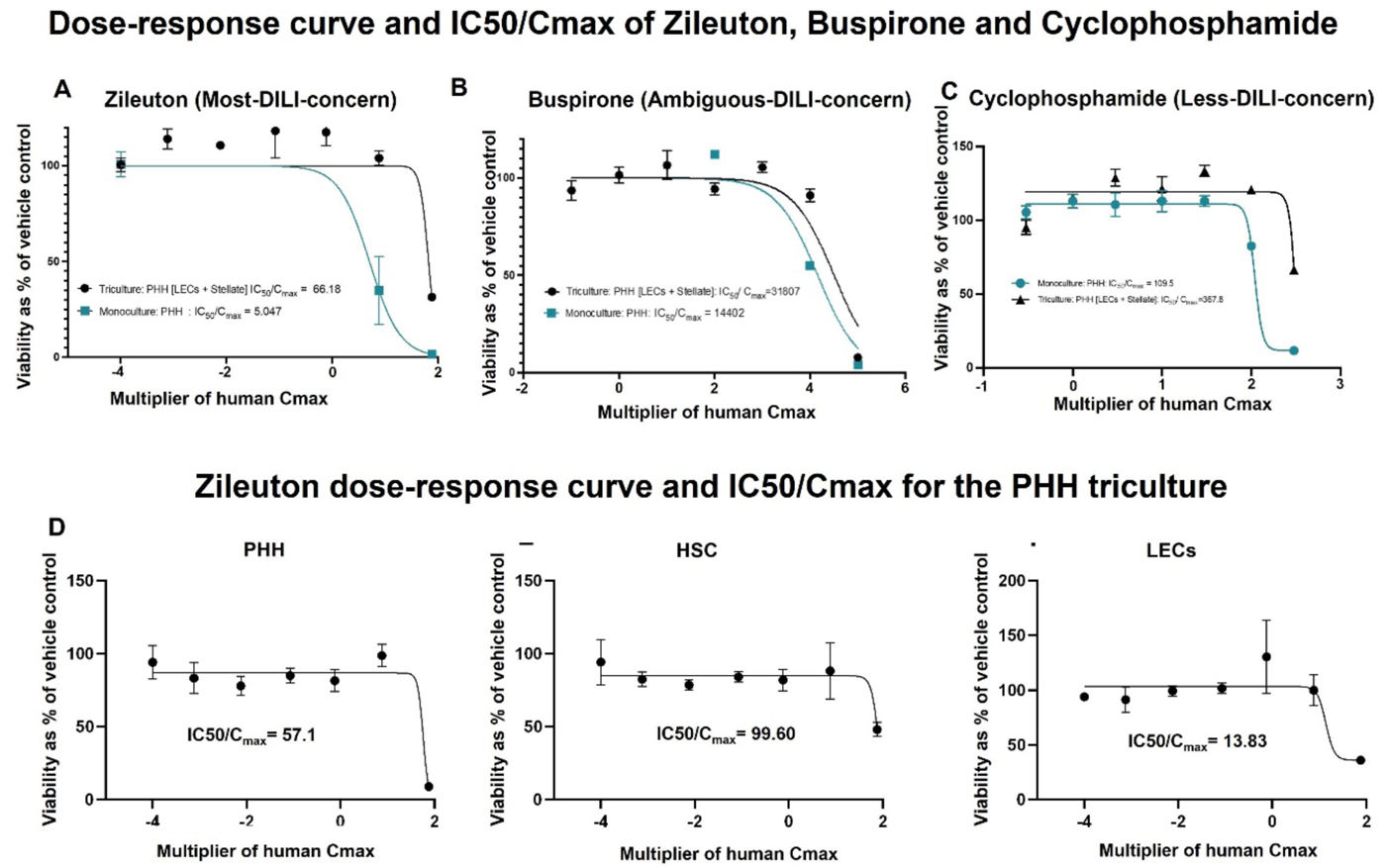
Toxicity evaluation of Zileuton, Buspirone, and Cyclophosphamide on monoculture and triculture models. Cell viability, reported on the y-axis in all graphs, is expressed as the percentage of ATP production of the vehicle control. Concentrations of the compounds, reported on the x-axis in all graphs, are expressed as multipliers of the human C_max_. The graphs show data points and dose-response curves obtained by applying a nonlinear regression. **A:** PHH viability associated with monoculture (blue) and triculture (black) models inclusive of fluid flow were treated with Zileuton at concentrations ranging between 0.0001 human C_max_ and 100 human C_max_. **B:** PHH viability associated with monoculture (blue) and triculture (black) models inclusive of fluid flow were treated with Busporine at concentrations ranging between 0.1 human C_max_ and 100000 human C_max_ **C:** PHH viability associated with monoculture (blue) and triculture (black) models inclusive of fluid flow were treated with Cyclophosphamide at concentrations ranging 0.3 human C_max_ and 300 human C_max_ (x axis). Cell viability (y axis) of PHH (**D**), HSC (**E**) and LECs (**F**) under static conditions in triculture, treated with Zileuton at concentrations ranging between 0.0001 human C_max_ and 100 human C_max_ (x axis). Each data point was obtained from n = 3 wells. Data are reported as mean ± SEM.

Overall, the addition of KC to the model did not markedly improve ATP, urea, albumin and CYP3A4 production relatively to the triculture model, whilst the same parameters were increased by the introduction of fluid flow in all models and across different timepoints. Therefore, triculture and monoculture models inclusive of fluid flow were used for toxicity screening experiments. In particular, the compounds tested were Zileuton (Fig. 7A), Buspirone (Fig. 7B) and Cyclophosphamide (Fig. 7C), respectively considered to be most-DILI-concern, ambiguous-DILI-concern and less-DILI-concern drugs by the U.S. FDA (58). Compounds testing was initiated at timepoints that were found to be associated with ascending trends in CYP3A4 production according to the results obtained from the characterisation experiment. In particular, the drug dosing protocol began on day 2 and day 4 for monocultures and tricultures respectively. Each dose of the drug was given every 24 h and the viability of PHH was analysed after a 7 days period. The IC50/C_max_ of Zileuton associated to the triculture was 66.18 mM, which was 1311.3% higher than the monoculture value of 5.047 mM, indicating that NPCs might exert a protective effect on PHH. A similar effect was observed with regards to Buspirone, as the IC50/C_max_ associated with tricultures was 31807 mM, 220.9% higher than the value associated to monocultures (14402 mM), and with regards to Cyclophosphamide, as the IC50/C_max_ associated with tricultures was 367.8 mM, 335.9% higher than the value obtained from monocultures (109.5 mM).

NS-based models can be disassembled and data can be acquired in relation to each individual NS, as shown in Figure 7, which depicts the viability of PHH (Fig. 7D), HSC (Fig. 7E) and LECs (Fig. 7F) in the triculture model upon treatment with Zileuton, in the static condition. In particular, LECs (IC50/C_max_ = 13.83) were more vulnerable to the toxic effects of Zileuton compared to PHH (IC50/C_max_ = 57.1) and HSC (IC50/C_max_ = 99.6).

## Discussion

Animal studies encounter significant limitations due to substantial differences in drug metabolism and pharmacokinetics between animals and humans (36, 37). A recent survey conducted in the pharmaceutical industry shed light on the limited concordance between preclinical liver toxicity findings and clinical outcomes (6). This finding is consistent with prior studies demonstrating the limited predictive capability of preclinical models for human liver toxicity (4). These studies underscore the limitations of current preclinical testing paradigms in predicting DILI in humans, particularly for compounds with poorly characterized dose-response relationships or unique mechanisms of toxicity. After the U.S. FDA released the Modernization Act 2.0, which allows the use of alternatives to animal testing to investigate the safety and effectiveness of a drug, a need for realistic human *in vitro* models of the liver for DILI screening has emerged (38). The main advantages of this approach will be the reduction of both time and costs of drug development, whilst following the principles of the “3 Rs” in relation to the replacement, reduction, and refinement of animal studies in the context of safety and efficacy assessments (39).

In this work, a novel platform called NANOSTACKS™ (NS) was utilized for the assembly of complex hepatic coculture models in a high-throughput 24-well plate format inclusive of fluid flow. The models included monocultures of PHH, tricultures of PHH, HSC and LECs, as well as tetracultures comprising PHH, HSC, LECs and KC. The models were characterized based on parameters associated with hepatic function, both in the absence and presence of fluid flow. The tricultures and monocultures incorporating fluid flow were further evaluated for their reliability in the context of DILI screening. This evaluation was conducted using three different compounds to assess the models’ effectiveness in predicting hepatic responses.

NS utilized for the development the complex models used in this work include a polycarbonate body and a transparent porous PET membrane, thus using materials that do not absorb tested drugs. On the other hand, a considerable number of devices used for the development of complex coculture models are fabricated using polydimethylsiloxane (PDMS), which absorb a wide spectrum of biochemical compounds, thus altering experimental outcomes of DILI screening (40–42). Additionally, both drug delivery challenges and the presence of a necrotic core are associated with the use of spheroids, due to their tightly assembled cellular geometry hampering the diffusion of nutrients and compounds to the innermost cellular layers. Conversely, *in vitro* models based on NS allow the unobstructed diffusion of compounds and nutrients towards all cellular layers (43). Moreover, whilst variability can be associated with the geometry of spheroids and organoids (43), NS-based models have high reproducibility due to the standardised dimensions of the individual NS. Other features of NS also include optical transparency and the possibility to include fluid flow in the model. Additionally, NS are compatible with SBS-standard 24-well plates and plate-reader-based assays, in addition to biochemical assays relying on the use of supernatant, thus making NS a user-friendly platform.

In the tricultures and tetracultures models, the ratios of PHH, LECs, HSCs, and KCs are congruent with those associated to native liver tissue (44). In particular, whilst most coculture models include a NPC:PHH ratio of 1:2 to 1:6 (16,45–46), in this study a 1:3 ratio was used, to increase the production of any paracrine signalling molecules associated to NPC.

Cytochrome P450 enzymes (CYPs), including CYP3A4, are crucial for the first-pass metabolism of xenobiotics. The inclusion of fluid flow increased CYP3A4 production relatively to the static condition at multiple timepoints. The CYP3A4 production observed in all NS-based human liver models was maintained throughout the entire duration of the study, as observed in MPS-based PHH models (47). Similarly to the results associated to CYP3A4 production, fluid flow also increased ATP production on multiple timepoints relatively to the static condition.

Albumin production was detectable across all three models, and was increased by the inclusion of fluid flow in most timepoints. In particular, in fluid flow-inclusive models, albumin secretion was found to be within the range associated with the *in vivo* human production rate (37-105 µg per day per 1 million hepatocytes) (39,48) on day 7 in monocultures, on day 11, 14 and 26 in tricultures, and on day 11 in tetracultures, indicating that these models effectively support hepatocyte functionality. Comparisons can be drawn with regards to other studies. In the work conducted by Tasnim et al., collagen sandwich cultures and spheroids including rat hepatocytes have shown relatively higher albumin levels compared to NS-based human-relevant models (49). However, in the same study, it was noted that rat hepatocytes-derived spheroids had higher albumin production compared to human-derived hepatic spheroids (49), and therefore the higher production rate associated to rat-derived models as compared to NS-based human models might be due to the hepatocytes origin rather than to the substrate onto which cells are cultured. Nonetheless, despite their albumin production, rat-derived hepatocytes cannot be considered to be as human-relevant as PHH (39). With regards to studies conducted on human cells, in the study by Rodriguez-Fernandez et al., PHH cultures on a 2D surface did not reach the human *in vivo* albumin production threshold (50) therefore indicating that PHH cultured on NS might adopt a more human-relevant phenotype as opposed to culture on standard well-plates. However, comparisons with more advanced culture systems including PHH spheroids lead to mixed conclusions. In the study conducted by Messner et al., spheroids developed using PHH and liver-derived NPC had higher albumin production rates than NS-based models, over a 5-week period (51). However, in the study conducted by Esch et al., long-term human liver organoid cultures over 14 days had a lower rate of albumin production compared to NS-based, fluid-flow inclusive models (51). Mixed results can be obtained with regards to comparisons with MPS-based models, with albumin production being relatively higher or lower than NS-based models depending on factors such as the timepoint, platform and PHH donor (14,47,53).

Urea production in all three NS-based, fluid flow-inclusive models was above the threshold of *in vivo* human urea production (56-159 µg per day per 1 million hepatocytes) value up to day 7 (39,54). Urea production in NS hepatic models in this study was higher compared to models based on rat-derived hepatocytes included in 2D monolayers (54), collagen sandwich cultures (48,55), spheroid models grown for 7 days (48,56) and micropatterned coculture of iPSC-derived human hepatocytes cultured for 28 days (31), demonstrating the importance of including human-derived primary cells in realistic *in vitro* models (52). As discussed in relation to albumin production, comparisons with studies analysing the urea production in MPS-based model hold mixed results depending on donor, platform and timepoint (14, 57). In summary, similarly to MPS platforms, NS-based PHH models inclusive of fluid flow can replicate human-relevant hepatic markers, such as *in vivo* albumin production, whilst being user-friendly and fitting into a 24-well plate format.

To evaluate the capacity of fluid flow-inclusive, NS-based monoculture and triculture models to be used as a screening platform for DILI, we examined the hepatotoxic effects of Zileuton, Buspirone and Cyclophosphamide. The PHH included in the triculture model consistently demonstrated greater resistance to cytotoxicity compared to the PHH in monoculture models, particularly in response to Zileuton and Cyclophosphamide. The enhanced PHH resistance to toxicity effects can be considered an advantageous feature in a DILI screening platform, as it could reduce the incidence of false positives and allow for more accurate estimations of human toxic dosages. Additionally, an advantageous feature of NS-based models is the possibility to analyse separately each NS upon drug testing, in order to assess the cell-specific cytotoxicity of a compound. In this work, this was demonstrated using the drug Zileuton. This differential toxicity analysis can be technically challenging in other types of coculture models, such as 2D mixed cocultures and spheroids.

In conclusion, NS can be used for the development of *in vitro* liver models that are compatible with plate-reader based assays. Multicellular human liver models developed using NS have shown fluid flow-dependent increases in the production of ATP, CYP3A4, albumin and urea. NS-based models can be used as DILI screening platforms, with PHH in triculture models exhibiting greater resistance to toxicity relatively to monocultures. Therefore, NS represents a promising tool for developing complex hepatic *in vitro* models with a view to reduce the high attrition rate associated with the drug development process. Future work will address the limitations of this research by expanding the donor pool, increasing the number of drugs tested, and evaluating their effects on liver functionality. Additionally, NS could also be used for the development of hepatic disease models, such as nonalcoholic steatohepatitis.

## ACKNOWLEDGEMENTS

We wish to thank Innovate UK for funding this project (Grant: 10035032). NANOSTACKS™ is a patented technology and consists of a family of patents including UK (Granted: GB1602146), US (Granted: US16/075136), and pending applications in Europe (EP1713365.9) and under the WIPO (PCT/GB2017/090286).

## References

1. Andrade RJ, Chalasani N, Björnsson ES, Suzuki A, Kullak-Ublick GA, Watkins PB, Devarbhavi H, Merz M, Lucena MI, Kaplowitz N, Aithal GP. Drug-induced liver injury. Nature Reviews Disease Primers. 2019 Aug 22;5(1):58.

2. Watkins PB. Drug safety sciences and the bottleneck in drug development. Clinical Pharmacology & Therapeutics. 2011 Jun;89(6):788–90.

3. Hornberg JJ, Laursen M, Brenden N, Persson M, Thougaard AV, Toft DB, Mow T. Exploratory toxicology as an integrated part of drug discovery. Part I: Why and how. Drug discovery today. 2014 Aug 1;19(8):1131–6.

4. Olson H, Betton G, Robinson D, Thomas K, Monro A, Kolaja G, Lilly P, Sanders J, Sipes G, Bracken W, Dorato M. Concordance of the toxicity of pharmaceuticals in humans and in animals. Regulatory toxicology and pharmacology. 2000 Aug 1;32(1):56–67.

5. McGill MR, Jaeschke H. Animal models of drug-induced liver injury. Biochimica et Biophysica Acta (BBA)-Molecular Basis of Disease. 2019 May 1;1865(5):1031–9.

6. Monticello TM, Jones TW, Dambach DM, Potter DM, Bolt MW, Liu M, Keller DA, Hart TK, Kadambi VJ. Current nonclinical testing paradigm enables safe entry to First-In-Human clinical trials: The IQ consortium nonclinical to clinical translational database. Toxicology and applied pharmacology. 2017 Nov 1;334:100–9.

7. Serras AS, Rodrigues JS, Cipriano M, Rodrigues AV, Oliveira NG, Miranda JP. A critical perspective on 3D liver models for drug metabolism and toxicology studies. Frontiers in cell and developmental biology. 2021 Feb 22;9:626805.

8. Han JJ. FDA Modernization Act 2.0 allows for alternatives to animal testing.

9. Collins SD, Yuen G, Tu T, Budzinska MA, Spring KJ, Bryant K, Shackel NA. In vitro models of the liver: disease modeling, drug discovery and clinical applications. Hepatocellular carcinoma. 2019:47–67.

10. Underhill GH, Khetani SR. Bioengineered liver models for drug testing and cell differentiation studies. Cellular and molecular gastroenterology and hepatology. 2018 Jan 1;5(3):426–39.

11. Proctor WR, Foster AJ, Vogt J, Summers C, Middleton B, Pilling MA, Shienson D, Kijanska M, Ströbel S, Kelm JM, Morgan P. Utility of spherical human liver microtissues for prediction of clinical drug-induced liver injury. Archives of toxicology. 2017 Aug;91:2849–63.

12. Takebe T, Sekine K, Enomura M, Koike H, Kimura M, Ogaeri T, Zhang RR, Ueno Y, Zheng YW, Koike N, Aoyama S. Vascularized and functional human liver from an iPSC-derived organ bud transplant. Nature. 2013 Jul 25;499(7459):481–4.

13. Palma E, Doornebal EJ, Chokshi S. Precision-cut liver slices: a versatile tool to advance liver research. Hepatology international. 2019 Jan 15;13:51–7.

14. Ewart L, Apostolou A, Briggs SA, Carman CV, Chaff JT, Heng AR, Jadalannagari S, Janardhanan J, Jang KJ, Joshipura SR, Kadam MM. Qualifying a human Liver-Chip for predictive toxicology: Performance assessment and economic implications. Biorxiv. 2021 Dec 16:2021–12.

15. Borgström A, Filippi BG, Hewitt P, Wolf A, Gebauer M. Expression Analysis of Exosomal MicroRNAs in Chlorpromazine Treated Human Hepatic Spheroids: A Promising Tool for Novel Drug-Induced Liver Injury Biomarker Discovery. Applied In Vitro Toxicology. 2023 Jun 1;9(2):65–76.

16. Bell CC, Chouhan B, Andersson LC, Andersson H, Dear JW, Williams DP, Söderberg M. Functionality of primary hepatic non-parenchymal cells in a 3D spheroid model and contribution to acetaminophen hepatotoxicity. Archives of toxicology. 2020 Apr;94:1251–63.

17. LeCluyse EL. Human hepatocyte culture systems for the in vitro evaluation of cytochrome P450 expression and regulation. European journal of pharmaceutical sciences. 2001 Jul 1;13(4):343–68.

18. Bell CC, Dankers AC, Lauschke VM, Sison-Young R, Jenkins R, Rowe C, Goldring CE, Park K, Regan SL, Walker T, Schofield C. Comparison of hepatic 2D sandwich cultures and 3D spheroids for long-term toxicity applications: a multicenter study. Toxicological Sciences. 2018 Apr 1;162(2):655–66.

19. Dash A, Inman W, Hoffmaster K, Sevidal S, Kelly J, Obach RS, Griffith LG, Tannenbaum SR. Liver tissue engineering in the evaluation of drug safety. Expert opinion on drug metabolism & toxicology. 2009 Oct 1;5(10):1159–74.

20. Uyama N, Shimahara Y, Kawada N, Seki S, Okuyama H, Iimuro Y, Yamaoka Y. Regulation of cultured rat hepatocyte proliferation by stellate cells. Journal of hepatology. 2002 May 1;36(5):590–9.

21. Thomas RJ, Bhandari R, Barrett DA, Bennett AJ, Fry JR, Powe D, Thomson BJ, Shakesheff KM. The effect of three-dimensional co-culture of hepatocytes and hepatic stellate cells on key hepatocyte functions in vitro. Cells Tissues Organs. 2006 Jul 4;181(2):67–79.

22. Kidambi S, Sheng L, Yarmush ML, Toner M, Lee I, Chan C. Patterned co-culture of primary hepatocytes and fibroblasts using polyelectrolyte multilayer templates. Macromolecular bioscience. 2007 Mar 8;7(3):344–53.

23. Domansky K, Inman W, Serdy J, Dash A, Lim MH, Griffith LG. Perfused multiwell plate for 3D liver tissue engineering. Lab on a Chip. 2010;10(1):51–8.

24. Chia SM, Lin PC, Yu H. TGF-β1 regulation in hepatocyte-NIH3T3 co-culture is important for the enhanced hepatocyte function in 3D microenvironment. Biotechnology and bioengineering. 2005 Mar 5;89(5):565–73.

25. Bhandari RN, Riccalton LA, Lewis AL, Fry JR, Hammond AH, Tendler SJ, Shakesheff KM. Liver tissue engineering: a role for co-culture systems in modifying hepatocyte function and viability. Tissue engineering. 2001 Jun 1;7(3):345–57.

26. Ohno M, Motojima K, Okano T, Taniguchi A. Up-regulation of drug-metabolizing enzyme genes in layered co-culture of a human liver cell line and endothelial cells. Tissue Engineering Part A. 2008 Nov 1;14(11):1861–9.

27. Chiew GG, Fu A, Perng Low K, Qian Luo K. Physical supports from liver cancer cells are essential for differentiation and remodeling of endothelial cells in a HepG2-HUVEC co-culture model. Scientific reports. 2015 Jun 8;5(1):10801.

28. Zinchenko YS, Schrum LW, Clemens M, Coger RN. Hepatocyte and kupffer cells co-cultured on micropatterned surfaces to optimize hepatocyte function. Tissue engineering. 2006 Apr 1;12(4):751–61.

29. Li F, Cao L, Parikh S, Zuo R. Three-dimensional spheroids with primary human liver cells and differential roles of Kupffer cells in drug-induced liver injury. Journal of pharmaceutical sciences. 2020 Jun 1;109(6):1912–23.

30. Han B, Mo H, Svarovskaia E, Mateo R. A primary human hepatocyte/hepatic stellate cell co-culture system for improved in vitro HBV replication. Virology. 2021 Jul 1;559:40–5.

31. Ware BR, Berger DR, Khetani SR. Prediction of drug-induced liver injury in micropatterned co-cultures containing iPSC-derived human hepatocytes. Toxicological sciences. 2015 Jun 1;145(2):252–62.

32. Chatterjee S, Richert L, Augustijns P, Annaert P. Hepatocyte-based in vitro model for assessment of drug-induced cholestasis. Toxicology and applied pharmacology. 2014 Jan 1;274(1):124–36.

33. Deharde D, Schneider C, Hiller T, Fischer N, Kegel V, Lübberstedt M, Freyer N, Hengstler JG, Andersson TB, Seehofer D, Pratschke J. Bile canaliculi formation and biliary transport in 3D sandwich-cultured hepatocytes in dependence of the extracellular matrix composition. Archives of toxicology. 2016 Oct;90:2497–511.

34. Saxton SH, Stevens KR. 2D and 3D liver models. Journal of Hepatology. 2023 Apr 1;78(4):873–5.

35. Driessen R, Zhao F, Hofmann S, Bouten C, Sahlgren C, Stassen O. Computational characterization of the dish-in-a-dish, a high yield culture platform for endothelial shear stress studies on the orbital shaker. Micromachines. 2020 May 29;11(6):552.

36. Lewis DF, Ioannides C, Parke DV. Cytochromes P450 and species differences in xenobiotic metabolism and activation of carcinogen. Environmental health perspectives. 1998 Oct;106(10):633–41.

37. Peter JO, Chan K, Silber PM. Human and animal hepatocytes in vitro with extrapolation in vivo. Chemico-biological interactions. 2004 Nov 1;150(1):97–114.

38. Adashi EY, O’Mahony DP, Cohen IG. The FDA modernization Act 2.0: drug testing in animals is rendered optional. The American Journal of Medicine. 2023 Sep 1;136(9):853–4.

39. Baudy AR, Otieno MA, Hewitt P, Gan J, Roth A, Keller D, Sura R, Van Vleet TR, Proctor WR. Liver microphysiological systems development guidelines for safety risk assessment in the pharmaceutical industry. Lab on a Chip. 2020;20(2):215–25.

40. Ma LD, Wang YT, Wang JR, Wu JL, Meng XS, Hu P, Mu X, Liang QL, Luo GA. Design and fabrication of a liver-on-a-chip platform for convenient, highly efficient, and safe in situ perfusion culture of 3D hepatic spheroids. Lab on a Chip. 2018;18(17):2547–62.

41. Carius P, Weinelt FA, Cantow C, Holstein M, Teitelbaum AM, Cui Y. Addressing the ADME Challenges of Compound Loss in a PDMS-Based Gut-on-Chip Microphysiological System. Pharmaceutics. 2024 Feb 20;16(3):296.

42. Van Meer BJ, de Vries H, Firth KS, van Weerd J, Tertoolen LG, Karperien HB, Jonkheijm P, Denning C, IJzerman AP, Mummery CL. Small molecule absorption by PDMS in the context of drug response bioassays. Biochemical and biophysical research communications. 2017 Jan 8;482(2):323–8.

43. Mehta G, Hsiao AY, Ingram M, Luker GD, Takayama S. Opportunities and challenges for use of tumor spheroids as models to test drug delivery and efficacy. Journal of controlled release. 2012 Dec 10;164(2):192–204.

44. Zeilinger K, Freyer N, Damm G, Seehofer D, Knöspel F. Cell sources for in vitro human liver cell culture models. Experimental Biology and Medicine. 2016 Sep;241(15):1684–98.

45. Hurrell T, Kastrinou-Lampou V, Fardellas A, Hendriks DF, Nordling Å, Johansson I, Baze A, Parmentier C, Richert L, Ingelman-Sundberg M. Human liver spheroids as a model to study aetiology and treatment of hepatic fibrosis. Cells. 2020 Apr 14;9(4):964.

46. Ware BR, Durham MJ, Monckton CP, Khetani SR. A cell culture platform to maintain long-term phenotype of primary human hepatocytes and endothelial cells. Cellular and molecular gastroenterology and hepatology. 2018 Jan 1;5(3):187–207.

47. Rubiano A, Indapurkar A, Yokosawa R, Miedzik A, Rosenzweig B, Arefin A, Moulin CM, Dame K, Hartman N, Volpe DA, Matta MK. Characterizing the reproducibility in using a liver microphysiological system for assaying drug toxicity, metabolism, and accumulation. Clinical and translational science. 2021 May;14(3):1049–61.

48. Ballmer PE, McNurlan MA, Milne E, Heys SD, Buchan V, Calder AG, Garlick PJ. Measurement of albumin synthesis in humans: a new approach employing stable isotopes. American Journal of Physiology-Endocrinology and Metabolism. 1990 Dec 1;259(6):E797–803.

49. Tasnim F, Singh NH, Tan EK, Xing J, Li H, Hissette S, Manesh S, Fulwood J, Gupta K, Ng CW, Xu S. Tethered primary hepatocyte spheroids on polystyrene multi-well plates for high-throughput drug safety testing. Scientific Reports. 2020 Mar 16;10(1):4768.

50. Rodriguez-Fernandez J, Garcia-Legler E, Villanueva-Badenas E, Donato MT, Gomez-Ribelles JL, Salmeron-Sanchez M, Gallego-Ferrer G, Tolosa L. Primary human hepatocytes-laden scaffolds for the treatment of acute liver failure. Biomaterials Advances. 2023 Oct 1;153:213576.

51. Messner S, Agarkova I, Moritz W, Kelm JM. Multi-cell type human liver microtissues for hepatotoxicity testing. Archives of toxicology. 2013 Jan;87(1):209–13.

52. Esch MB, Prot JM, Wang YI, Miller P, Llamas-Vidales JR, Naughton BA, Applegate DR, Shuler ML. Multi-cellular 3D human primary liver cell culture elevates metabolic activity under fluidic flow. Lab on a Chip. 2015;15(10):2269–77.

53. Török E, Lutgehetmann M, Bierwolf J, Melbeck S, Düllmann J, Nashan B, Ma PX, Pollok JM. Primary human hepatocytes on biodegradable poly (l-lactic acid) matrices: A promising model for improving transplantation efficiency with tissue engineering. Liver transplantation. 2011 Feb;17(2):104–14.

54. Rudman D, DiFulco TJ, Galambos JT, Smith RB, Salam AA, Warren WD. Maximal rates of excretion and synthesis of urea in normal and cirrhotic subjects. The Journal of clinical investigation. 1973 Sep 1;52(9):2241–9.

55. Bale SS, Golberg I, Jindal R, McCarty WJ, Luitje M, Hegde M, Bhushan A, Usta OB, Yarmush ML. Long-term coculture strategies for primary hepatocytes and liver sinusoidal endothelial cells. Tissue Engineering Part C: Methods. 2015 Apr 1;21(4):413–22.

56. Kim MK, Jeong W, Jeon S, Kang HW. 3D bioprinting of dECM-incorporated hepatocyte spheroid for simultaneous promotion of cell-cell and-ECM interactions. Frontiers in Bioengineering and Biotechnology. 2023 Nov 13;11:1305023.

57. Xiao RR, Lv T, Tu X, Li P, Wang T, Dong H, Tu P, Ai X. An integrated biomimetic array chip for establishment of collagen-based 3D primary human hepatocyte model for prediction of clinical drug-induced liver injury. Biotechnology and bioengineering. 2021 Dec;118(12):4687–98.

58. Chen M, Suzuki A, Thakkar S, Yu K, Hu C, Tong W. DILIrank: the largest reference drug list ranked by the risk for developing drug-induced liver injury in humans. Drug discovery today. 2016 Apr 1;21(4):648–53.

